# A neural network model for the orbitofrontal cortex and task space acquisition during reinforcement learning

**DOI:** 10.1101/116608

**Authors:** Zhewei Zhang, Zhenbo Cheng, Zhongqiao Lin, Chechang Nie, Tianming Yang

## Abstract

Reinforcement learning has been widely used in explaining animal behavior. In reinforcement learning, the agent learns the value of the states in the task, collectively constituting the task state space, and use the knowledge to choose actions and acquire desired outcomes. It has been proposed that the orbitofrontal cortex (OFC) encodes the task state space during reinforcement learning. However, it is not well understood how the OFC acquires and stores task state information. Here, we propose a neural network model based on reservoir computing. Reservoir networks exhibit heterogeneous and dynamic activity patterns that are suitable to encode task states. The information can be extracted by a linear readout trained with reinforcement learning. We demonstrate how the network acquires and stores the task structures. The network exhibits reinforcement learning behavior and its aspects resemble experimental findings of the OFC. Our study provides a theoretical explanation of how the OFC may contribute to reinforcement learning and a new approach to understanding the neural mechanism underlying reinforcement learning.

## Introduction

Even the simplest reinforcement learning (RL) algorithm captures the essence of operant conditioning in psychology and animal learning (Rescorla & Wagner, 1972). That is, actions that are rewarded tend to be repeated more frequently; actions that are punished are more likely to be avoided. RL requires one to understand the structures of the task and evaluate the value of the states in the task state space. Several studies have investigated the possible brain structures that may be involved in RL (Daw, Gershman, Seymour, Dayan, & Dolan, 2011; Glascher, Daw, Dayan, & O'Doherty, 2010; Haber, Kim, Mailly, & Calzavara, 2006; Kennerley, Behrens, & Wallis, 2011; Schultz, Dayan, & Montague, 1997). Notably, the orbitofrontal cortex (OFC) has been hypothesized to represent the task space and encode task states (Wilson, Takahashi, Schoenbaum, & Niv, 2014). Several lesion studies showed that the animals with OFC lesions exhibited deficits acquiring task information for building a task structure (Hornak et al., 2004; Izquierdo, Suda, & Murray, 2004; Takahashi et al., 2011). On the other hand, experiments that recorded single unit activities in the OFC have mostly found that the OFC neurons encode many aspects of reward information, including reward value (J. L. Jones et al., 2012; Padoa-Schioppa, 2011; Padoa-Schioppa & Assad, 2006; Rudebeck, Mitz, Chacko, & Murray, 2013; Wallis & Miller, 2003), probability (Kennerley & Wallis, 2009), risk (O'Neill & Schultz, 2015), information value (Blanchard, Hayden, & Bromberg-Martin, 2015), abstract rules (Wallis, Anderson, & Miller, 2001), and strategies (Tsujimoto, Genovesio, & Wise, 2011). Yet, it is not obviously how these recording experiment results can be reconciled with the hypotheses that the OFC is involved in representing task structures. There is a lack of theory how task structures themselves may be encoded and represented by a neural network, and what sort of neuronal firing properties we expect to find in neurophysiological experiments. Furthermore, we do not know how to teach a task-agnostic neural network to acquire the structure of the task just based on trial and error.

In the current study, we provide a solution based on the reservoir network (Buonomano & Maass, 2009; Laje & Buonomano, 2013; Maass, Natschlager, & Markram, 2002). Reservoir networks are recurrent networks with fixed connections. Within a reservoir network, neurons are randomly and sparsely connected. Importantly, the internal states of a reservoir exhibit rich temporal dynamics, which represents a nonlinear transformation of its input history and can be very useful for encoding task state sequences. The information encoded by the network can be extracted with a linear output, which can be trained during learning. Reservoir networks have been shown to exhibit dynamics similar to that observed in the prefrontal cortex (Barak, Sussillo, Romo, Tsodyks, & Abbott, 2013; Cheng, Deng, Hu, Zhang, & Yang, 2015; Enel, Procyk, Quilodran, & Dominey, 2016).

One key feature of our reservoir-based network model that makes learning task structures possible is to include reward itself as an input to the reservoir. Thereby, the network dynamics represents a combination of not only the sensory events, but also the reward outcome. The reinforcement learning helps to shape the output of the reservoir, essentially picking out the action that will lead to the desired event sequences that lead to rewards.

We demonstrate with two commonly used learning paradigms how the network model works. Task event sequences, including reward events, are provided as inputs to the network. A simple yet biologically feasible reward-dependent Hebbian learning algorithm is used to adjust its output weights. We show that our network model can solve problems with different task structures and reproduce behavior experiments previously conducted in animals and humans. We further demonstrate the similarities between the reservoir network and the OFC. Manipulations to our network reproduce the behavior of animals with OFC lesions. Moreover, the reservoir neurons’ response patterns resemble characteristics of the OFC neurons reported from previous electrophysiological experiments. Taken together, these results suggest a simple mechanism that naturally leads to the acquisition of task structure and therefore supports RL. Finally, we propose some future experiments that may be used to test our model.

## Results

We describe our results in three parts. We start with using our network to model a classical reversal learning task. We take advantage of the simplicity of the task to explain the principal ideas behind the network model and why we think the network resembles the OFC. Then we show such a network may be applied to more complex scenarios in which the OFC has been shown to play important roles. Finally, to further illustrate the similarities between our network model and the OFC, we demonstrate how the selectivity of the neurons in the network may resemble experimental findings in the OFC during value-based decision making.

### Reversal Learning

In a classical reversal learning task, the animals have to keep track of the reward contingency of two choice options that may be reversed during a test session (Izquierdo et al., 2004; B. Jones & Mishkin, 1972). Normal animals were found to learn reversals faster and faster, which has been used as an indication of their ability of learning the structure of the task (Wilson et al., 2014). Such behavior was however found to be impaired in animals with OFC lesions or with lesions that contained fibers passing near the OFC (Izquierdo et al., 2004; Rudebeck, Saunders, Prescott, Chau, & Murray, 2013). These animals were not able to learn reversals faster and faster when they were repeatedly tested. The learning impairments could be explained by a deficit in acquiring and representing the task structure (Wilson et al., 2014).

Our neural network model consists of a state encoding layer (SEL), which is a reservoir network. It receives three inputs and generates two outputs (Fig 1a). The three inputs to the SEL are the two choice options *A* and *B*, together with a reward input that indicates whether the choice yields a reward or not in the current trial. The outputs represent choice actions *A* and *B* for the next trial. We use the neural activity of the SEL at the end of the input to determine the SEL’s output.

**Figure 1.**
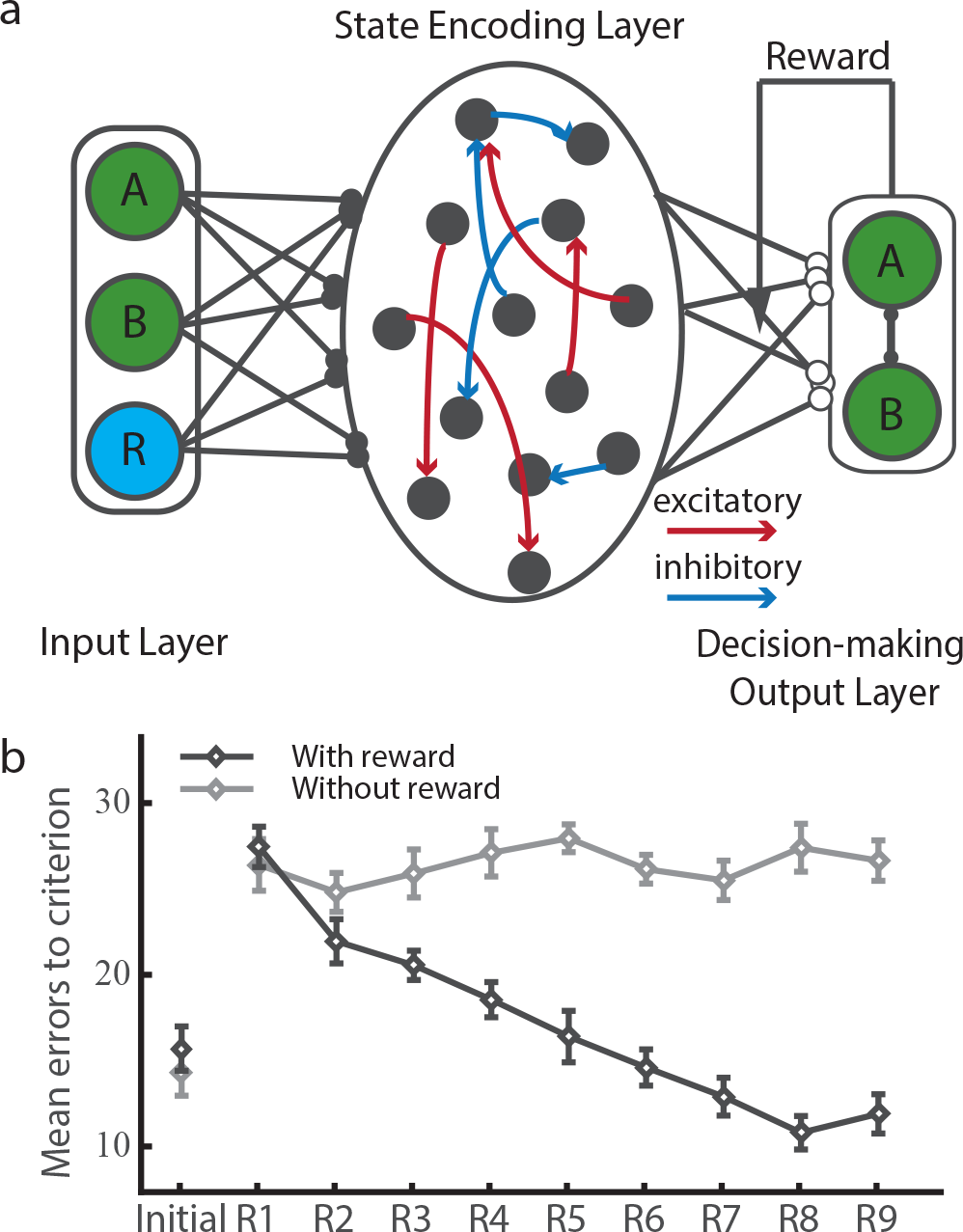
(a) The schematic diagram of the model. The network is composed of three parts: input layer (IL), the state encoding layer (SEL) and the decision-making output layer (DML).(b) The number of the error trials made before the network achieves the performance threshold. The dark line indicates the performance of the network with the reward input; the light line indicates the performance of the network without the reward input as a model for animals of OFC lesions.

The network is able to reproduce animals’ behavior. The number of error trials that takes for the network to achieve the performance threshold, which is set at 93% in the initial learning and at 80% in the subsequent reversals, decreases as the network goes through more and more reversals (Fig 1b). Interestingly, a learning deficit similar to that found in OFC-lesion animals is observed if we remove the reward input to the SEL (Fig 1b). As the OFC and its neighboring brain areas such as the ventromedial prefrontal cortex (vmPFC) are known to receive both the sensory inputs and reward inputs from sensory and reward circuitry in the brain, removing the reward input from our model mimics the situation where the brain has to learn without functioning structures in or near the OFC.

Neurons in the SEL, as expected from a typical reservoir network, show highly heterogeneous response patterns. Some neurons are found to encode the stimulus identity, some neurons encode reward, and others show mixed tuning (Fig 2a). A principal component analysis (PCA) based on the population activity shows that the network can distinguish all four possible task states: choice *A* rewarded, choice *A* not rewarded, choice *B* rewarded, and choice *B* not rewarded (Fig 2b).

**Figure 2.**
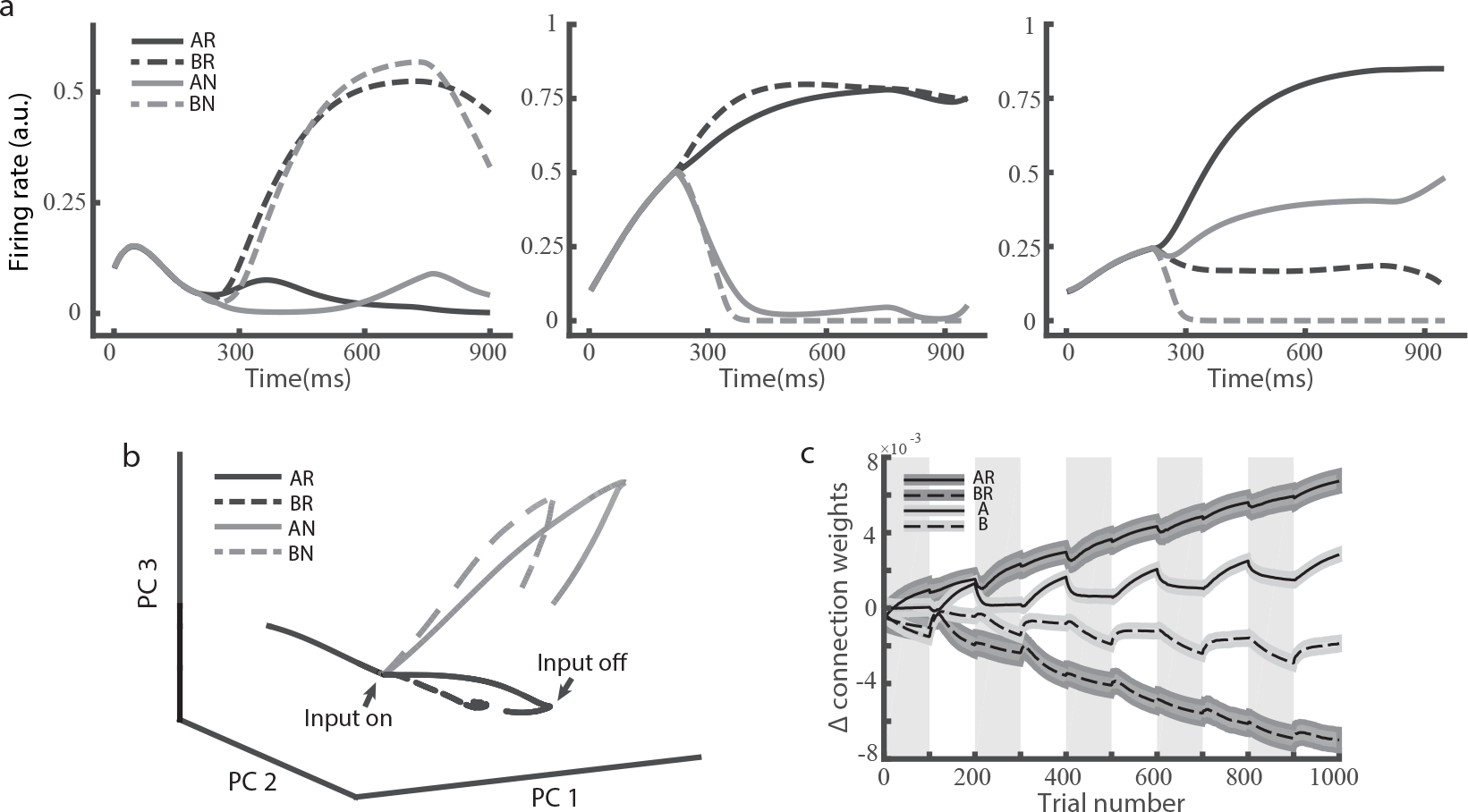
(a) Selectivity of three example neurons in the reservoir network. Input units are set to 1 from 200ms to 700ms. Left panel: an example neuron that encodes choice options; middle panel: an example neuron that encodes reward outcomes; right panel: an example neuron with mixed selectivity. (b) PCA on the network population activity. The network states are plotted in the space spanned by the first 3 PCA components. The activities in different conditions are differentiated after the cue onset. (c) The difference between the connection weights between SEL neurons and the DML unit *A* and DML unit *B*. The SEL neurons are grouped according to their selectivities. For example, *AR* represents the group of neurons that respond most strongly when the input units *A* and *R* are both activated. The gray and white area indicates the blocks in which the option *A* and the option *B* leads to the reward, respectively.

The ability to distinguish these states is essential for learning. To understand the task acquisition behavior exhibited by our model, we study how neurons with different selectivity contribute to the learning (Fig 2c). We find that readout weights of the neurons that are selective to the combination of stimulus and reward inputs (e.g. *AR* and *BR*) are mostly affected by the learning. The difference between the weights of their connections to the outputs *A* and *B* keeps growing despite repeated reversals. In contrast, the weights of the output connections of pure stimulus-selective neurons only wiggle around the baseline between reversals.

The difference between these two groups of neurons explains why our network achieves flexible learning behavior only when the reward input is available. Let us first consider the *AR* neurons, which are selective for the situation when choice *A* leads to reward. In these *A*-rewarded blocks, the connections between the *AR* neurons and the DML neuron of choice *A* are strengthened. When the reward contingency is reversed and now choice *A* leads to no reward, the connections between the *AR* neurons and choice *A* are not affected very much. That is because the group of *AN* neurons instead of the *AR* neurons are activated in the blocks when choice *A* is not rewarded. As the result, the connections between the *AN* neurons and the DML neuron of choice *B* are strengthened and the connections between the *AN* neurons and the DML neuron of choice *A* are weakened. When the reward contingency is flipped again, the connections between the *AR* neurons and the DML neuron of choice *A* are strengthened further. This way, the learning is never erased by the reversals, and the network learns faster and faster. In comparison, let us now consider the *A* neurons, which encode only the sensory inputs and are activated whenever input *A* is present. In the *A*-rewarded blocks, the connections between the *A* neurons and the DML neuron of choice *A* are strengthened. In *B*-rewarded blocks, the connections between the *A* neurons and the DML neuron of choice *A* are however weakened when the network chooses *A* and gets no reward, and the learning in the previous block is reversed. Thus, the output connections of *A* neurons only fluctuate around the baseline with the reversals. They do not contribute much to the learning, and the overall behavior of the network is mostly driven by neurons that are activated by the combination of reward input and sensory inputs. Removing *R* deactivates these neurons and leads to the structure agnostic behavior.

It is important to note that although the reinforcement learning algorithm employs the same small learning rate for both the intact network and the “OFC-lesion” network, the former only requires a few number of trials to acquire a reversal in the later stage of training, indicating learning may not have to be slow with a small learning rate.

### Two-stage Markov decision task

We further test our network with a two-stage decision making task. The task is similar to the Markov decision task used previously in several human fMRI studies and used to study the model-based reinforcement learning behavior in humans (Glascher et al., 2010). In this task, the subjects have to choose between two options *A1* and *A2*. Their choices then lead to two intermediate outcomes *B1* and *B2* at different but fixed probabilities. The choice of *A1* more likely leads to *B1*, and the choice of *A2* is more likely followed by *B2*. Importantly, the final reward is contingent only on these intermediate outcomes, and the contingency is reversed across blocks (Fig 3a). Thus, the probability of getting a reward is higher for *B1* in one block and becomes lower in the next block. The probabilistic association between the initial choices and the intermediate outcomes never changes. The subjects are not informed of the structure of the task, and they have to figure out the best option by tracking not only the reward outcomes but also the intermediate outcomes.

**Figure 3.**
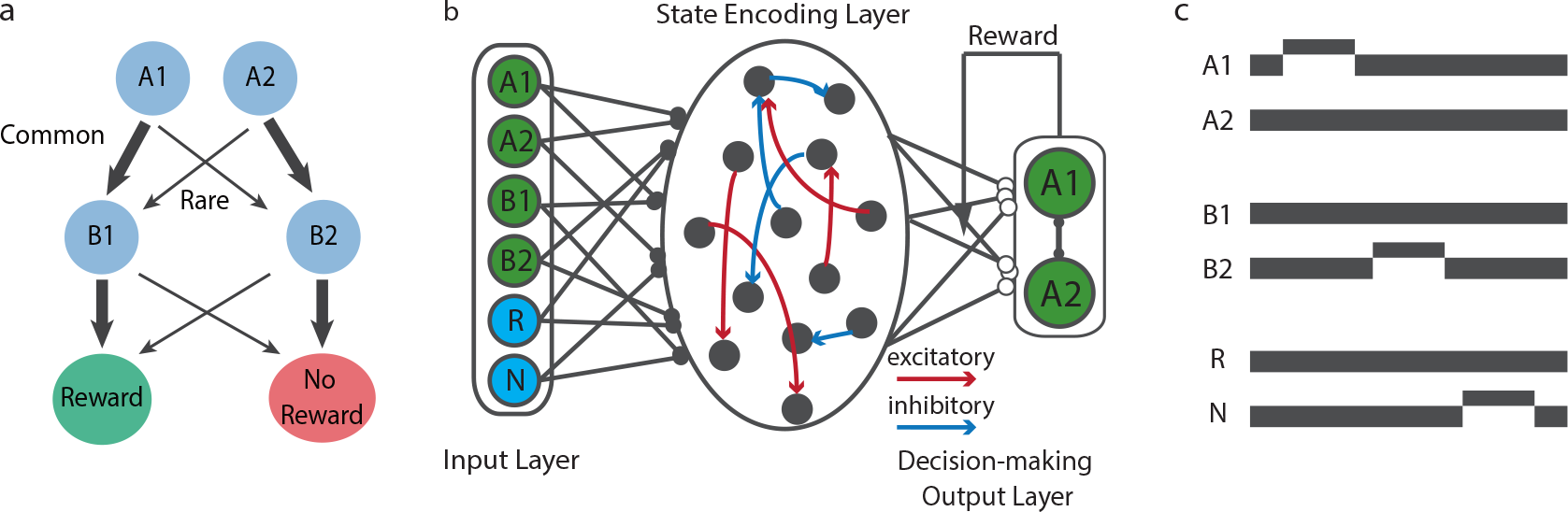
(a) Task structure of the two-stage Markov decision task. Two options *A1* and *A2* are available, they lead to two intermediate outcomes *B1* and *B2* at different probabilities. The width of the arrows indicates the transition probability. Intermediate outcomes *B1* and *B2* lead to rewards at different probability, but the reward contingency of the intermediate outcomes is reversed between blocks. (b) The schematic diagram of the model. It is similar to Fig 1a. The only difference is that there are more input units. (c) Units in the input layer are activated sequentially. In the example trial, option *A1* is chosen, *B2* is presented, and no reward is obtained.

We keep our network model mostly the same as in the previous task. Here, we have two additional input units that reflect the intermediate outcomes (Fig 3b). To demonstrate our network model’s capability of encoding sequential events, the input units are activated sequentially in our simulations as they are in the real experiment (Fig 3c). We also add a *non-reward* input unit whose activity is set to 1 when a reward is not obtained at the end of a trial. The additional non-reward input facilitates learning but does not change the results qualitatively.

For a simple temporal difference learning strategy without using any knowledge of task structure, the probability of repeating the previous choice only depends on the reward outcome. The probability of repeating the previous choice is higher when a reward is obtained than when no reward is obtained. The intermediate outcome is ignored. However, this is no longer the case when the task structure is taken into account. For example, consider the situation when the subject initially chooses *A1*, the intermediate outcome happens to be *B2*, and a reward is obtained. If the subject understands *B2* is an unlikely outcome of choice *A1* (rare), but a likely outcome of choice *A2* (common), a reward obtained after the rare event *B2* should actually motivate the subject to switch from the previous choice and choose *A2* the next time. The subject should always choose the option that is more likely to lead to the intermediate outcome that is currently associated with the better reward.

To quantify the learning behavior, we first evaluate the impact of the previous trial’s outcome on the current trial. We classify all trial outcomes into four categories: common-rewarded (*CR*), common-unrewarded (*CN*), rare-rewarded (*RR*) and rare-unrewarded (*RN*). Here, common and rare indicate whether the intermediate outcome is the more likely outcome of the chosen option or not. Glascher et al (Glascher et al., 2010) showed that the model based learning led to a higher probability of repeating the previous choice in the *CR* and *RN* conditions. This is also what we observe in our network model’s behavior (Fig 4a).

**Figure 4.**
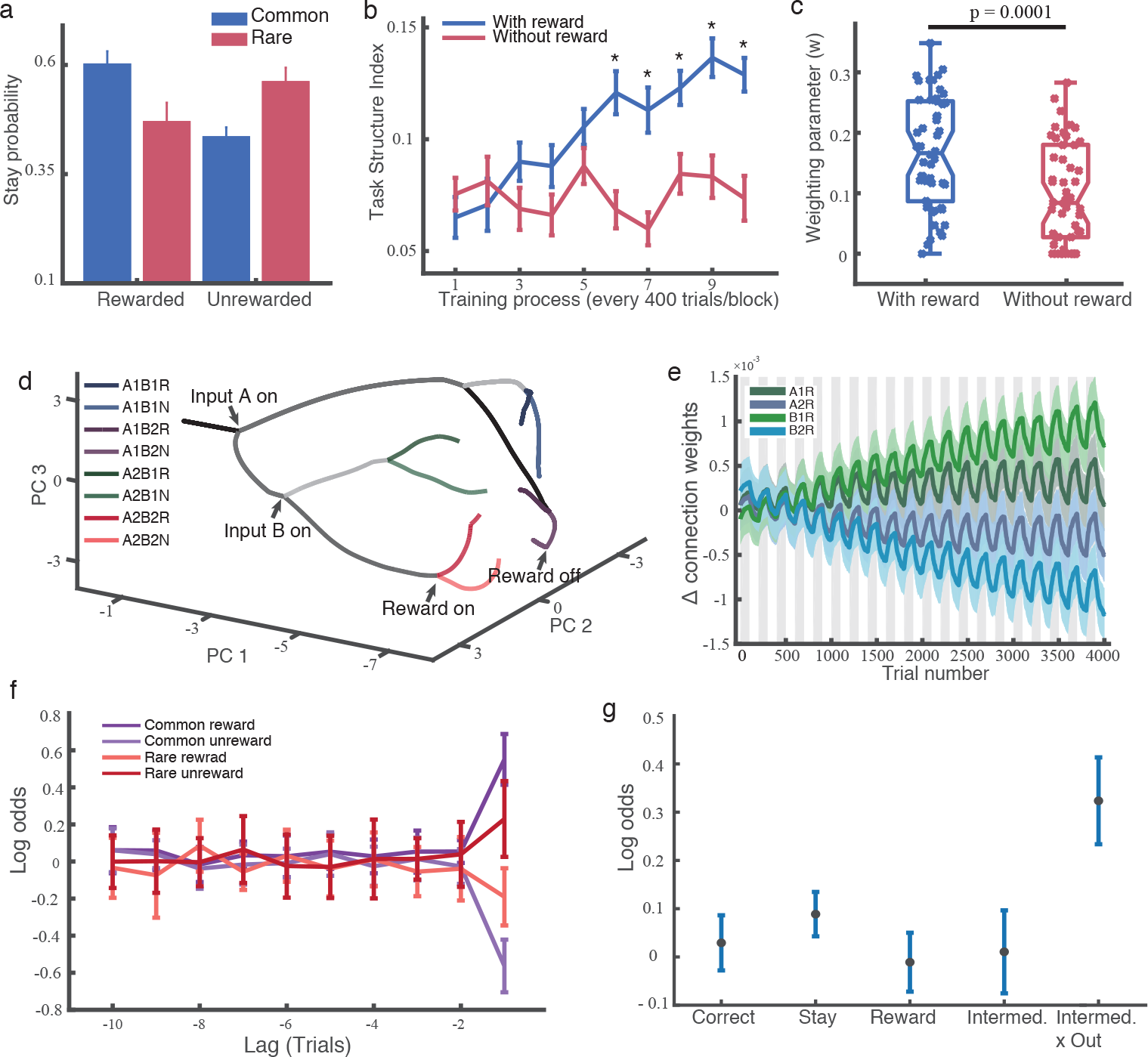
(a) Factorial analysis of choice behavior. The agent is more likely to repeat the choice under the conditions common-rewarded (*CR*) and rare-unrewarded (*RN*) than under the conditions common-unrewarded (*CU*) and rare-rewarded (*RR*). (b) The task structure index keeps growing in the intact network (blue line), but stays at a low level when the network is without its reward input (red line). (c) Fitting the behavioral performance with a mixture of task-agnostic and task-aware algorithms. The weight parameter *w* for learning with the knowledge of the task structure is significantly larger for the intact network (blue data points) than the network without the reward input (red data points). Each data point represents a simulation run. (d) PCA on the network population activity. The network states are plotted in the space spanned by the first 3 PCA components. The network can distinguish all 8 different states. (e) The weight differences between the connections between SEL neurons and the DML unit *A1* and DML unit *A2*. Similar to Fig 2c. The gray and white areas indicate the blocks in which intermediate outcome *B1* is more likely to lead to a reward and the blocks in which *B2* is more likely to lead to a reward, respectively. (f) Logistic regression shows that only the last trial’s state affect the choice. The regression includes four different states (intermediate outcome x reward outcome) for each trial up to 10 trials before the current trials. Error bars show SEM across simulation runs. (g) Logistic regression reveals that only the combination of the intermediate states and the reward outcome in the last trial affects the decision. The factors being evaluated are: Correct—a tendency to choose the better choice in current block; Reward—a tendency to repeat the previous choice if it is rewarded; Stay—a tendency to repeat the previous choice; Intermed.—a tendency to repeat the same choice following common intermediate outcomes; Intermed. x Out–a tendency to repeat the same choice dependent on the interaction between intermediate outcomes (common/rare) and reward outcome.

To illustrate how the network acquires the task structure, we define the task-structure index, which represents the tendency of employing task structure information (see the *Method*). The task-structure index grows larger as the training goes on (Fig 4b). It indicates that the network learns the structure of the task gradually and transits to a more efficient behavior from an initially task-agnostic behavior. Similar to our findings in the first task, the SEL without the reward input does not show this transition (Fig 4b). We further quantify the contribution of task structure information to the network behavior using a model fitting procedure previously described by Glascher et al. (Glascher et al., 2010), and the network without the reward input shows a significantly smaller weight for the usage of task structure, suggesting it is worse at picking up the task structure (Fig 4c).

Again, a PCA on the SEL population activity shows that the SEL distinguishes different task states (Fig 4d). Because of the structure of the task in which the contingency between the first stage options and the intermediate outcomes is fixed, the network only needs to find out the current reward contingency of the intermediate outcomes. We found that the learning picks out the most relevant neurons that encode the contingency between the intermediate outcomes and the reward outcomes (*B1R*, *B2R*, etc.). Their connection weights to the DML neurons show better and better differentiation of the two choices throughout the training (Fig 4e). In contrast, the connection weights of the neurons that encode the association between the first stage options and the reward outcomes (*A1R*, *A2R*, etc.) are less differentiated.

These results suggest that the network acquires the task structure. It understands the contingency between the intermediate outcomes and the reward outcomes is the key to the performance. Thus, its choice only depends on the interaction between the intermediate outcome and the reward outcome of the last trial, but not on the other factors (Fig 4f and 4g). The network behavior is similar to the *Reward-as-cue agent* described by Akam et al. (Akam, Costa, & Dayan, 2015).

### Value representation by the OFC

Previous electrophysiology studies have shown that OFC neurons encode value during economic choices (Padoa-Schioppa & Assad, 2006; Wallis & Miller, 2003). Among these value-encoding neurons, studies have identified multiple classes of neurons encoding a variety of information, including the value of individual offers (offer value), the value of the chosen option (chosen value), and the identity of the chosen option (chosen identity) (Cai & Padoa-Schioppa, 2014; Padoa-Schioppa, 2013).

Here we show that our network model may explain this apparent heterogeneous value encoding in the OFC. Here we model a two-alternative economic choice task by providing two inputs to the SEL, representing the value of each option (Fig 5a). The network model can reproduce the choice behavior of monkeys (Fig 5b) (Padoa-Schioppa & Assad, 2006). Then we study the selectivity of the SEL neurons. We find not only neurons that encode the value of each option (offer value neurons, middle panel in Fig 6a), but also neurons that encode the value of the chosen option (chosen value neurons, left panel in Fig 6a). Furthermore, a proportion of neurons show the selectivity for the choice as previously reported (chosen identity neurons, right panel in Fig 6a). We classify the neurons in the reservoir network into 10 categories as described in Padoa-Schioppa and Assad (Padoa-Schioppa & Assad, 2006). Interestingly, we are able to find neurons in 9 of the 10 categories (Fig 6b, c). The only missing category (neurons encoding other/chosen value) was also very rare in the experimental data. Although the proportions of neurons encoding each category are not an exact copy of the experimental data, but the similarity is apparent. This is surprising given that we do not tune the internal connections of the SEL to the task. The heterogeneity is naturally expected from a reservoir network, but it takes much more effort to explain with recurrent network models that have a well-defined structure (Daie, Goldman, & Aksay, 2015; Rustichini & Padoa-Schioppa, 2015).

**Figure 5.**
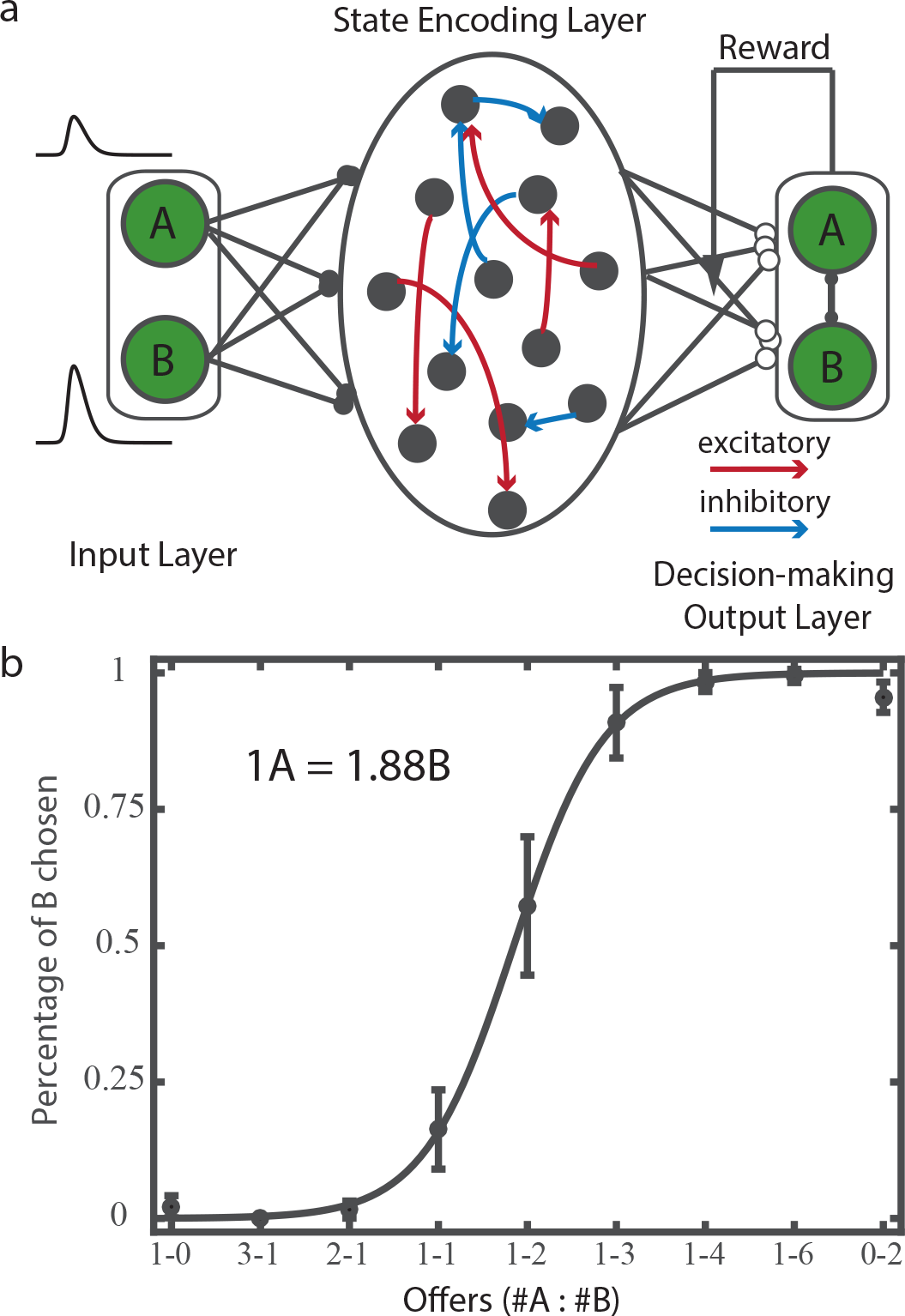
(a) The schematic diagram of the model. The input neurons’ responses are not a step function as in the previous paradigms, illustrated in the left side of the panel. (b) Choice pattern. The relative value preference calculated based on the network behavior is indicated on the top left, and the actual relative value preference used in the simulation is 1*A* = 2*B*.

**Figure 6.**
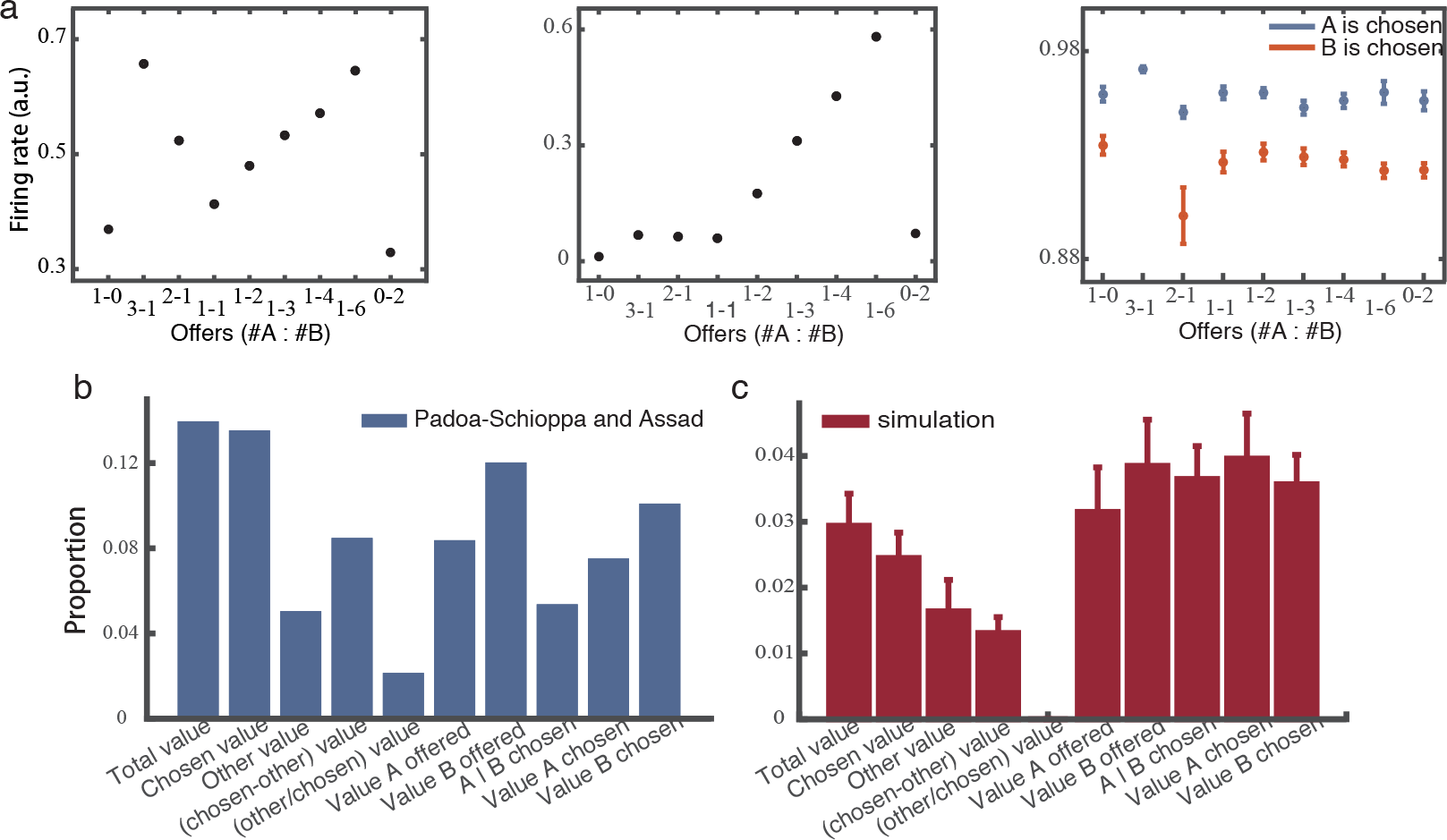
(a) Three example neurons in the SEL. Left panel: a neuron that encodes chosen value; middle panel: a neuron that encodes offer value; right panel: a neuron that encodes chosen juice. (b) The proportions of the neurons with different selectivities from a previous experimental study (Padoa-Schioppa & Assad, 2006). (c) The proportions of the neurons in our reservoir network with different selectivities.

## Discussion

So far, we have shown that a simple reservoir-based network model may acquire task structures. The more interesting question is that why the network is capable of doing so and how this network model may help us to understand the functions of the OFC.

### Encoding of the task space

We place a reservoir network as the centerpiece of our model. Reservoir networks are large, distributed, nonlinear dynamical recurrent neural networks with fixed weights. Because of recurrent networks’ complicated dynamics, they are especially useful in modeling temporal sequences including languages (Rodriguez, 2001; Suykens, Vandewalle, & Moor, 1996). They have been shown to be Turing equivalent (Kilian & Siegelmann, 1996) and capable of approximating arbitrary dynamical systems (Funahashi & Nakamura, 1993). In our model, the reservoir network encodes the combinations of inputs that constitute the task state space. States are encoded by the activities of the reservoir neurons, and the learned action values are represented by the weights of the readout connections.

There are several reasons why we choose reservoir networks to construct our model. First reason is that we would like to pair our network model with reinforcement learning. Reservoir networks have fixed internal connections; the training occurs only at the readout. The number of parameters is thus much smaller, which could be important for efficient reinforcement learning. Generality is another benefit offered by reservoir networks. Because the internal connections are fixed, we can use the same network to solve a different problem by just training a different readout. The reservoir can be seen as a general-purpose task state representation network. Lastly, our results as well as several other studies show that neurons in reservoir networks – although with untrained connections weights – show properties similar to that observed in the real brain (Barak et al., 2013; Cheng et al., 2015; Sussillo & Abbott, 2009), suggesting local plasticity may not play a role as important as previously thought.

The fact that the internal connections are fixed in a reservoir network means that the selectivity of the reservoir neurons is also fixed. This may seem at odds with the experimental findings of many OFC neurons shifting their encodings rapidly during reversals (Rolls, Critchley, Mason, & Wakeman, 1996). However, these observations may be interpreted differently. The neurons that were found to have different responses during reversals might be in fact encoding rewards. On the other hand, there is evidence that OFC neurons with inflexible encodings during reversals might be more important for flexible behavior (Schoenbaum, Saddoris, & Stalnaker, 2007).

The choice of a reservoir network as the center piece of task event encoding may appear questionable to some. We do not train the network to learn task event sequences. Instead, we use the dynamic patterns elicited by task event sequences as bases for learning. This approach has obvious weaknesses. One is that the chaotic nature of network dynamics limits how well the task states can be encoded in the network. However, the purpose of our network model is not to solve complicated tasks. Instead, we would like to argue this is a more biologically-realistic model than many other recurrent networks. First, it does not depend on supervised learning to learn task event sequences (Song, Yang, & Wang, 2017; Sussillo & Abbott, 2009). Second, although the network performance may appear to be very limited by task complexity, the real brain, however, also has limited capacity in learning multi-stage tasks (Akam et al., 2015). Lastly, we show that a reservoir network can describe OFC neuronal responses during value-based decision making. Several other studies have also shown that reservoir networks may be a useful model of the prefrontal cortex (Barak et al., 2013; Cheng et al., 2015).

### Reward input to the reservoir

One key observation is that reward events must also be provided as inputs to the reservoir layer for the network model to perform well. Including reward events allows the network to establish associations between sensory stimuli and rewards, thus facilitates task structure acquisition. Although reward modulates neural activities almost everywhere in the cortex, the OFC is unique in its role of encoding the association between sensory stimuli and rewards. Removing the reward input to the reservoir mimics the situation when animals cannot rely on such an association to learn tasks. In this case, the reservoir is still perfectly functional in terms of encoding task events other than rewards. This is similar to the situation when animals have to depend on their other memory structures in the brain –such as hippocampus or other medial temporal lobe structures – for learning. The importance of the reward input to the reservoir explains the key role that the OFC plays in RL.

Several recent studies reported that selective lesions in the OFC did not reproduce the behavior deficits in reversal learning previously seen if the fibers passing through or near the OFC were spared (Rudebeck, Saunders, et al., 2013). Since these fibers probably carry the reward information from the midbrain areas, these results do not undermine the importance of reward inputs. Presumably, when the lesion is limited to the OFC, the projections that carrying the reward information were still available to or might even be redirected to other neighboring prefrontal structures, including ventromedial prefrontal cortex, which might take over the role of the OFC and contribute to the learning in animals with selective OFC lesions.

### Model-based Reinforcement Learning

The acquisition of task structure is a prerequisite for model-based learning. Therefore, it is interesting to ask whether our network model is able to achieve model-based learning. The two-stage task that we model has been used in human literature to study model-based learning (Daw et al., 2011; Glascher et al., 2010). Our model, although exhibiting behavior similar to human subjects, can be categorized as the Reward-as-cue agent that was described and categorized as a form of model-free reinforcement learning agent by Akam et al. (Akam et al., 2015). However, it is conceivable that our network could support model-based learning by providing the task structure to the downstream network layers.

### Extending the network

The performance of our network depends on several factors. First, it is important that reservoir should be able to distinguish between different task states. The number of possible task states may be only 4 or 8 as in our examples, or may be impossibly large even if the number of inputs increases only slightly. The latter is due to the infamous combinatorial explosion problem. One may alleviate the problem by introducing learning in the reservoir to weed out irrelevant combinations. Second, the dynamics of the reservoir should allow information to be maintained long enough until the decision is made. The recent developed gated recurrent neural networks may provide a solution with units that may maintain information for long periods (Chung, Gulcehre, Cho, & Bengio, 2014). Third, the model exhibits substantial variability between runs, suggesting the initialization may impact its performance. Further investigation is needed to make the model more robust. Last, we show that a reinforcement learning algorithm is capable of solving the relatively simple tasks in this study. However, it has been shown that reinforcement learning is in general not very efficient for extracting information from reservoir networks. A possible solution is to introduce additional layers to help with the readout (Cheng et al., 2015).

### Testable Predictions

Our model makes several testable predictions. First, because of the reservoir structure, the inputs from the same source should be represented evenly in the network. For example, in a visual task, different visual stimuli should be represented at roughly the same strength in the OFC, even if their visual salience may be drastically different. Second, we should be able to find neurons encoding all relevant task parameters in the network. Third, reducing the number of inputs may make the network to be more efficient in certain tasks. This may seem counter-intuitive. But removing inputs reduces the number of states that the network has to encode, thus improves learning efficiency for tasks that do not require those additional states. For example, if we remove the reward input to the SEL, which is essential for learning tasks with volatile rewards, the network should however be more efficient at learning tasks in a more stable environment. Indeed, animals with OFC lesions were found to perform better than control animals when reward history was not important (Riceberg & Shapiro, 2012).

### Summary

Our network does not intend to be a complete model of how the OFC works. Instead of creating a complete neural network solution of reinforcement learning or the OFC, which is improbable at the moment, we are aiming at the modest goal of providing a proof of concept that approaches the critical problem of how the brain acquires the task structure with a biologically realistic neural network model. By demonstrating the network’s similarity to the experimental findings in the OFC, our study opens up new possibilities in future investigation.

## Materials and Methods

### Neural Network Model

The model is composed of three layers: an input layer (IL), a state encoding layer (SEL), and a decision-making output layer (DML) (Fig. 1a).

The units in the input layer represent the identities of sensory stimuli and the reward obtained.

The input neurons are sparsely connected to the SEL units. The connection weights 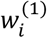 are set to 0 at a probability of *p*_IR_. Nonzero weights are assigned independently from a standard uniform distribution [0, 1].

In the SEL, there are *N*= 500 neurons. The neurons in the SEL are connected with a low probability *p*=0.1 and the connections are randomly and independently set from a Gaussian distribution with zero mean and a variance of *g*^2^/(*p*N)*, where the gain *g* acts as the control parameter in the SEL. Connections in the SEL could be both positive and negative.

Each neuron in the SEL is described by an activation variable *x_i_* for *i* = 1, 2, …, *N*, which is initialized with a normal distribution *N*(0, *σ*_ini_^2^) at the beginning of each trial. *x_i_* is updated at each time step (*dt* = 1ms) as follows:

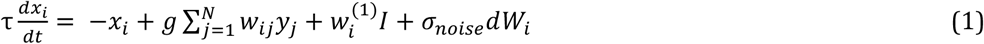

where *τ* represents the time constant, *w_ij_* is the synaptic weight between neurons *i* and *j*, *dW_i_* stands for the white noise, and *σ*_noise_ is its variance. The firing rate *y_i_* of neuron *i* is a function of the activation variable *x_i_* relative to a minimal firing rate *y*_min_=0 and the maximal rate *y*_max_=1:

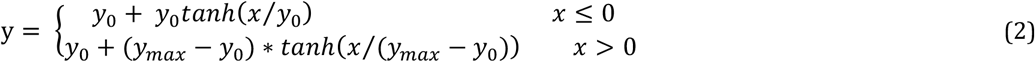

Here *y*_0_ = 0.1 is the baseline firing rate.

The SEL neurons project to the DML. The two competing neurons in the DML represent the two choices respectively. The total input of neuron *k* in the DML is

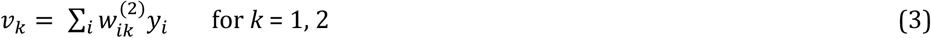

where *w_ik_^(2)^* is the weight of the synapse between neuron *i* in the SEL circuit and neuron *k* in the DML. The synaptic weights between the SEL and DML are randomly initialized with uniform distribution [0, 1], and normalized to keep the squared sum of synaptic weights projecting to the same DML unit equal to 1.

The synaptic weights between the SEL and DML are updated based on the choice and the reward outcome during the training phase. The stochastic choice behavior of our model is described by a softmax function:

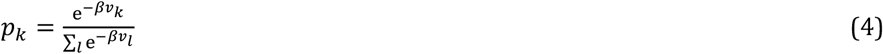

where *p_k_* represents the probability for choosing the choice *a_k,_* and the other choice is chosen with probability 1-*p_k_*. *β* adjusts the competition strength of two choices, and *v_k_* is the input of the DML unit *k*. The firing rate of the unit *k*, *y_k_*, is set to 1 if choice *a_k_* is chosen, otherwise it is set to 0.

### Reinforcement Learning

At the end of each trial, the weights between the SEL and the DML neurons are updated. The plastic weights in eq (3) in trial *n*+1 are updated as follows:

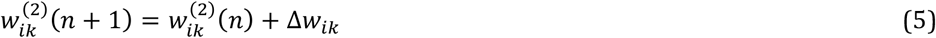

The update term Δ*w_ik_* depends on the reward prediction error and the responses of the neurons in the SEL circuit and DML:

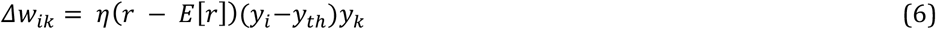

where *η* is the learning rate, and *r* is the reward. *E*[*r*] denotes the expected value. When the reward *r* is larger than E[r], the connections between the SEL neurons whose firing rate is above the threshold *y_th_* and the neurons in the DML would be strengthened, and the connections between the neurons whose firing rate is below *y_th_* and the neurons in the DML would be weakened. After each update, the weights 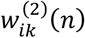 are normalized:

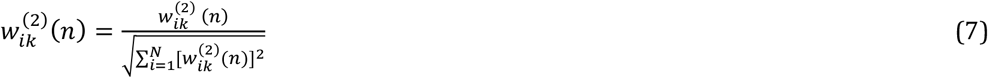

so that the vector length of 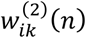remains constant. The normalization stops the weights from growing infinitely (Royer & Pare, 2003).

### Behavior Task

#### Reversal learning

The network has to choose between two options. One option leads to a reward, and the other does not. The stimulus-reward contingency is reversed every 100 trials. The criterion for learning is set to 28 correct trials in 30 successive trials for the initial learning and 24 correct trials in 30 successive trials for subsequent reversals.

The input layer units represent the identities of the two options and the reward. An option unit’s response is set to 1 for if the corresponding option is chosen in the current trial, otherwise it is set to 0. The reward unit’s response is set to 1 if the choice is rewarded in the current trial. The output of the network indicates its choice for the next trial. The network parameters are set as follows. Time constant *τ* = 100ms, network gain *g*=2, training threshold *y_th_* = 0.4, temperature parameter *β* = 4, learning rate η = 0.001, noise gain *σ*_noise_ =0.01, initial noise gain *σ*_ini_ = 0.01, input connection probability *p*_IR_=0.2.

The selectivity of neurons in the SEL is determined at the time point when the decision is made. A unit is defined as selective to a certain input or a combination of inputs if its responses are significantly higher under the condition when the input or all inputs of the combination are set to 1 than when some of them are set to 0.

#### Two-stage Markov decision task

The network has to make a choice between options *A1* and *A2*. *A1* leads to intermediate outcome *B1* at the probability of 80%, and *B2* at the probability of 20%. Vice versa, option *A2* leads to *B2* at the probability of 80%, and *B1* at a lower probability of 20%. The contingency between options (*A1*, *A2*) and intermediate outcomes (*B1*, *B2*) is fixed. Initially, *B1* leads to a reward at the probability of 80% and *B2* leads to reward at the probability of 20%. The reward contingency is reversed every 50 trials.

The input layer contains 6 units, representing the identities of two first stage options *A1* and *A2*, two intermediate outcomes *B1* and *B2*, and the reward and non-reward conditions, respectively. The activity of option unit *A1* or *A2* is set to 1 when the respective option is chosen. The activity of intermediate outcome unit *B1* or *B2* is set to 1 when the respective intermediate outcome is presented. The reward unit’s activity is set to 1 when a reward is obtained, and the non-reward unit’s activity is set to 1 when no reward is obtained. The units are activated sequentially, reflecting the sequential nature of the task. The *A* units are activated between 200 and 700ms after a trial starts, the *B* units between 700 and 1200ms, and the reward units between 1200 and 1700ms.

The output of the network indicates its choice. The network parameters are set as follows. Time constant *τ* = 500ms, Network gain *g*=2.25, training threshold *y_th_* = 0.2, temperature parameter *β* = 2, learning rate η = 0.001, noise gain *σ*_noise_ =0.01, initial noise gain *σ*_ini_ = 0.01, input connection probability *p*_IR_=0.2.

The selectivity of neurons in the SEL is determined at the time point when the decision is made. There are 8 conditions in this task, namely *A1B1R*, *A1B1N*, *A2B1R*, *A2B1N*, *A1B2R*, *A1B2N*, *A2B2R*, and *A2B2N*. For example, *A1B1R* indicates the condition when *A1* is chosen, intermediate outcome *B1* is presented, and a reward is obtained. A neuron’s preferred condition is the condition under which its activity is the largest and significantly higher than its activity under any other conditions. Then the neurons are grouped into different categories based on their preferred conditions. The neurons in category *A1R* are the neurons whose preferred conditions are *A1B1R*, *A1B2R*, *A2B1N* and *A2B2N*. All the preferred conditions of the neurons in category *A1R* provide evidence for associating *A1* with the reward. Similarly, the preferred conditions of the neurons in the category *B1N* are *A1B1N*, *A1B2R*, *A2B1N* and *A2B2R*. They provide evidence that *B1* is not associated with the reward.

##### Model fitting

In order to test how well the network uses the task structure information, we fit our data based on the model introduced by Daw et al. (Daw et al., 2011). The model fits the behavioral results with a mixture of model-free (task-agnostic) and model-based (task-aware) learning algorithm. In our simplified task, the network makes only one choice in each trial. The inverse temperature parameters *β_1_* is set to 2, which is also used to produce simulated behavioral choices. The parameter *p*, which captures the tendency for perseveration and switching, is set to 0, although all conclusions still hold when *p* is allowed to vary. The free parameters relevant in our task are *α_1_*, *α_2_*, *λ* and *w*. *α_1_* and *α_2_* are the learning rates in the learning algorithms with and without using task structure information, respectively. The eligibility *λ* represents how large proportion of credit from the reward can be given to the first states and actions in our task paradigm. *w* is the weight for how task information is employed. When *w* equals 1, the behavior uses complete task information. When *w* equals 0, the behavior is purely task agnostic. The fitting is done by a maximum likelihood estimation procedure.

##### Task-structure index

Inspired by the factorial analysis from Daw et al. (Daw et al., 2011), we define a task-structure (TS) index (eq.8) to quantify the tendency of repeating the choice in the last trial under different situations. The combination of the two reward outcomes and the two intermediate outcomes, common and rare, gives us four possible outcomes: common-rewarded (*CR*), common-unrewarded (*CN*), rare-rewarded (*RR*) and rare-unrewarded (*RN*). When the task structure is known, the agent is more likely to repeat the previous choice if the last trial is a *CR* or an *RN* trial. Higher TS index means that the behavioral pattern takes into account more task structure information.

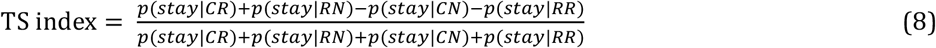

##### Task State Analysis

The task state analysis is similar to what was used by Akam et al. (Akam et al., 2015). Briefly, a logistic regression is used to estimate how states of past trials influence the current choice. The regression includes four different states (2 intermediate outcomes x 2 reward outcomes) for each trial up to 10 trials before the current trials.

Another logistic regression is used to estimate how several other potentially relevant factors affect choices. The factors considers include: Correct—a tendency to choose the better choice in current block; Reward—a tendency to repeat the previous choice if it is rewarded; Stay—a tendency to repeat the previous choice; Intermed.—a tendency to repeat the same choice following common intermediate outcomes; Intermed. x Out–a tendency to repeat the same choice dependent on the interaction between intermediate outcomes (common/rare) and reward outcome.

### Value-based economic choice task

Unlike the two previous paradigms, both options in this paradigm lead to a reward. Two input units represent the rewards associated with the two options, respectively. The input strength is proportional to reward magnitude. In our simulations, the reward *A* is valued twice as much as reward *B* for the same reward magnitude. The relative value preference between the two options is not provided as an input to the network directly, but used in calculating the expected value. The value of the reward is defined as the product of the relative value and the reward magnitude.

The activity of the input unit, f(*t*), is described by the following equations (Rustichini & Padoa-Schioppa, 2015).

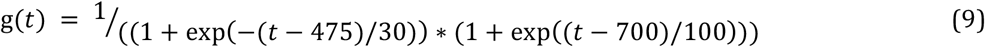

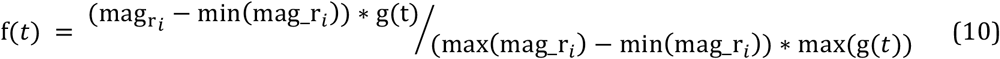

Here *t* is the time in the unit of ms within a trial, magr_*i*_is the magnitude of the reward type *i*in each trial, max (mag_r*i*_) is the maximal reward magnitude of reward type *i* within the block, and min (mag_r*_i_*)represents the minimal reward magnitude of reward type *i*, which is always 0 in our simulations. The expected value is the sum of the product of the probability of choosing the option and corresponding reward magnitude.

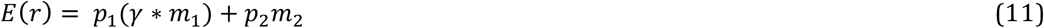

where *p_i_* and *m_i_* are the probability of choosing option *i* and its reward magnitude, and γ=2 is the relative value preference between the two reward options. Only the data from the trials after 6000 trials training are included for the analyses. The network parameters are set as follows. Time constant *τ* = 100ms, Network gain *g*=2.5, training threshold *y_th_* = 0.2, temperature parameter *β* = 4, learning rate ?? = 0.005, noise gain *σ*_noise_ =0.05, initial noise gain *σ*_ini_ = 0.2, input connection probability *p*_IR_=0.2.

As in Padoa-schioppa and Assad (Padoa-Schioppa & Assad, 2006), the following variables are defined for further analysis: total value (the sum of the value of two options), chosen value (the value of the chosen option), other value (the value of the unchosen option), value difference (chosen-other value), value ratio (other/chosen value), offer value (the value of the one option), chosen juice (the identity of the chosen option), and value A chosen (the value of the option A when option A is chosen).

We use an analysis similar to that in Padoa-schioppa and Assad (Padoa-Schioppa & Assad, 2006) to study the selectivity of SEL units during the post-offer period (0-500ms after the stimulus onset). Linear regressions are applied to each variable to fit the neural responses in this time window for each SEL unit separately. A variable is considered to explain the response of a neuron if the slope of a fitting linear function is significantly different from zero.

## Acknowledgements

Acknowledgements: This work is supported by the CAS Hundreds of Talents Program and Science and Technology Commission of Shanghai Municipality (15JC1400104) to T. Y., and by Public Projects of Zhejiang Province (2016C31G2020069) and the 3^rd^ Level in Zhejiang Province "151 talents project” to Z. C. We thank Yu Shan and Xiao-Jing Wang for discussions and comments during the study. The authors declare no competing financial or nonfinancial interests.

